# Pre-transplant TCR Network Topology Predicts Kidney Allograft Rejection Independent of HLA Mismatch

**DOI:** 10.64898/2026.05.04.722749

**Authors:** Nicholas Borcherding, Jes M. Sanders, Greg R. Martens, Naoka Murakami, Natnael Doilicho, Barbara L. Banbury, Jie He, Joseph R. Leventhal, James M. Mathew

## Abstract

Despite extensive pretransplant serological screening and HLA matching, 10–15% of kidney allografts experience acute rejection within the first year. Currently, risk stratification for transplantation relies primarily on antibody reactivity to HLA molecules, with no assessment of the T cell compartment before or after transplantation. In our previously established longitudinal cohort of 54 patients, T cell receptor β (TCRβ) sequencing was performed on paired kidney biopsy and peripheral blood samples. Here, we further analyzed the data to construct a comprehensive set of sequence-similarity networks and quantify over 30 network metrics. After adjusting for repertoire size, graft status was the strongest signal for the underlying differences in network metrics. Individuals who rejected the kidney graft generally exhibited more fragmented and less connected networks at baseline, with fewer interconnect T cell clones and more isolated sequences. Notably, pre-transplant peripheral blood mononuclear cell (PBMC) network topology alone predicted non-stable outcomes with an area under the curve (AUC) of 0.81, sensitivity of 76%, and specificity of 76%. The performance of this prediction model was independent of HLA mismatch, while changes in network topology at three months post-transplantation further improved prediction to an AUC of 0.88 (permutation p = 0.009). Collectively, TCR sequencing and network analysis represent a potential novel, non-invasive approach for pre-transplant risk stratification and immune monitoring, capturing functional immunological risk that may not be accessible through HLA genotyping or serology.

## Introduction

Kidney transplantation is the definitive treatment for end-stage renal disease. Despite progress in donor-recipient matching to minimize immunological risk, acute rejection occurs in 10-15% of recipients within the first year.^1^ The current paradigm in risk stratification relies on HLA matching and characterization of serological reactivity across donor-recipient pairs, the latter serving as an indirect low-resolution readout of the B cell receptor (BCR) repertoire alloreactivity. However, these tools are imperfect, rejection occurs in well-matched pairs^2^, while stable outcomes are observed despite high degrees of mismatch or serological reactivity. Moreover, non-HLA genetic incompatibilities between donor and recipient also contribute to allograft outcomes, suggesting that factors beyond HLA compatibility play a role in immunological tolerance.^3,4^ These lapses in prediction may in part be due to the absence of profiling a key component of the alloreactive response: T cells.

Through the somatic recombination of V(D)J gene segments, the T cell receptor (TCR) repertoire encodes the history and potential of the adaptive immune system. Bulk TCRβ sequencing now captures tens of thousands of clonotypes per sample, and a growing body of work has applied repertoire profiling to solid organ transplantation. In a systematic review of 37 studies spanning 2010–2021, Wong et al. found that decreased diversity is a universal feature of transplant recipients relative to healthy controls, but that the differentiating signal between patients with and without rejection lies in clonal expansion patterns rather than gross diversity changes.^5^ In the largest study to date (n=200), Sigdel et al. found that pre-transplant T-cell fraction and post-transplant repertoire turnover distinguished rejectors from stable patients, yet notably, clonality itself did not, suggesting that diversity metrics capture only part of the immunological signal.^6^ A subset of these studies used mixed lymphocyte culture to define alloreactive TCR fingerprints in donor-recipient pairings.^7–12^ Clone-tracking approaches offer greater specificity with two recent studies identifying persistent CD8+ donor-reactive clones despite induction therapy^10,11^, but require donor cells and multi-day culture, limiting clinical scalability. Critically, all of these approaches rely on scalar diversity metrics or individual clone identification.^5^ A notable exception of applied graph theory in transplant has shown that donor reactive clones exhibit increased modularity compared to the overall repertoire when examining the most expanded clones.^9^ Emerging evidence across autoimmunity, infection, and cancer immunology demonstrates that clonal relationships and structural organization carry predictive information beyond composition alone.^13–15^ This gap motivates a fundamentally different analytic framework, one that preserves the relational architecture of the repertoire rather than reducing it to summary statistics.

Sequence-similarity networks offer a framework that preserves the relational structure of the TCR repertoire.^13,15–17^ Rather than simply counting clonotypes, this approach maps the relationships between them. The concept is analogous to a social network: just as people (nodes) are connected by shared interests (edges), individual clonotypes (nodes) are linked by shared structural similarity (edges) when their sequences fall below a defined similarity threshold. From this representation, graph-theoretic metrics, including connectivity (the overall density of sequence-sharing across the repertoire), modularity (the degree to which the network segregates into distinct subgroups), and community structure (the identity and composition of those clusters), capture topological features that are invisible to scalar diversity indices. These properties may reflect the immunological forces shaping repertoire architecture: convergent selection pressures that drive sequence similarity among antigen-specific clones, patterns of clonal expansion that alter local network density, and the overall “readiness” of the repertoire to mount a coordinated response against specific antigens. We hypothesized that pre-transplant TCR network topology reflects the functional state of the recipient’s immune system in ways that predict allograft outcomes, and that this structural information is orthogonal to the risk captured by conventional HLA-based stratification.

Here, we apply sequence-similarity network analysis to a longitudinal TCRβ sequencing cohort of 54 kidney transplant recipients sampled across four timepoints (pre-transplant through 12 months post-transplant) in both peripheral blood and kidney biopsies. We constructed 297 TCR networks and computed over 30 graph-theoretic metrics, capturing connectivity, community structure, and global topology, with rigorous size normalization to address the known confounding of network metrics by repertoire size. We integrated these topological features with donor-reactive T cell annotations derived from mixed lymphocyte culture and clinical HLA mismatch data to evaluate whether network architecture provides predictive information independent of conventional risk stratification. We demonstrate that pre-transplant TCR network topology alone predicts graft outcomes, this signal is orthogonal to HLA mismatch, and that incorporating early post-transplant longitudinal dynamics further improves classification, establishing TCR network analysis as a potential novel framework for non-invasive, pre-transplant immunological risk assessment.

## Methods

### TCR Sequence Processing

TCRβ sequences were imported from the Adaptive Biotechnologies (Seattle, Washington) tab-delimited output, excluding nonproductive rearrangements (sequenceStatus ≠ “In”). When multiple gene assignments were present, *vGeneNameTies* and *jGeneNameTies* were used to fill missing entries in *vGeneName* and *jGeneName*, respectively. Gene names were simplified to their corresponding gene families by removing allele- and subgene-level suffixes (e.g., *TRBV12-03* to *TRBV12*) and correcting nonstandard nomenclature (e.g., *TCRBV* to *TRBV*). Leading zeroes were removed from single-digit gene numbers (e.g., *TRBV01* to *TRBV1*) to ensure consistency across annotations. For sequences with multiple gene assignments, all unique simplified gene families were retained and collapsed into a comma-separated list. Sequences with complementarity-determining region 3 (CDR3) with non-standard amino acid or lengths outside 5-25 residues were excluded from downstream analysis.

### Identification of Donor Reactive T Cell Clones (DRTC)

DRTC were identified as described previously.^11^ Briefly, DRTC were identified in the pretransplant setting using an anti-donor mixed lymphocyte reaction assay: Carboxyfluorescein diacetate succinimidyl ester (CFSE)-diluting CD4^+^ and CD8^+^ DRTC were flow sorted, and the TCRβ sequences were identified using TCRβ sequencing with the same processing steps as described above. DRTC were then tracked in post-transplant kidney biopsies and PBMC samples.

### Network Construction

Edit-distance-based similarity networks were constructed within strata defined by patient, timepoint, and tissue. For each stratum, clonotypes were aggregated across all library files mapped to that stratum, and duplicate clone pairs (defined by matching Vgene and CDR3 amino acid sequences or V, CDR3-AA) were removed. Networks were generated using the immApex (v1.4.4) R package, with CDR3 amino-acid sequences as nodes and undirected edges connecting sequences within a relative edit distance of 0.3 or less, based on normalized Levenshtein edit distance. To minimize spurious connectivity across V families, edges were permitted only between clonotypes sharing the same V gene (*filter.v* = TRUE). Node metadata (TRBV assignment) was retained for each edge to enable precise matching of donor-reactive labels. For each patient, distinct (V, CDR3-AA) pairs were retained for CD4 donor-reactive T cell clones (DRTC) and CD8 DRTC. DRTC annotations were integrated into each stratum network by exact matching on clones (V, CDR3-AA).

### Network Measurements

For each stratum network, we computed a comprehensive set of graph-level metrics using igraph (v2.2.1) R package. Basic structural properties included node count, edge count, density, and the fraction of nodes in the giant component. Degree distribution was characterized by mean degree, degree variance, coefficient of variation, and Shannon entropy. Weighted metrics based on edge similarity (defined as 1 minus the normalized edit distance) included node strength, weighted eigenvector centrality, and weighted local clustering coefficient. Community structure was detected using the Louvain algorithm, with optimal resolution determined by normalized mutual information and a grid search of k ∈ {0.5, 0.75, 1.0, 1.25, 1.5, 2}, yielding modularity and community count. K-core decomposition provided the maximum and mean coreness, as well as the fraction of nodes in the maximum core. For networks with fewer than 5,000 nodes, we also computed the mean shortest-path length and the diameter within the giant component.

DRTC-specific network metrics comprised the fraction of nodes annotated as CD4- or CD8-donor-reactive T cells, the internal edge density of DRTC-only subgraphs, DRTC assortativity (the tendency of DRTC clonotypes to connect preferentially with other DRTC clonotypes), and mixing matrix proportions. Community-level DRTC enrichment was evaluated using a hypergeometric test. To control for non-alloimmune antigen-experienced clonotypes, nodes were also annotated with known EBV and CMV specificities from the VDJdb database^18^ (minimum confidence score ≥ 1) using exact V-gene and CDR3 amino acid matching, and parallel viral-specific graph metrics were computed. Temporal metrics, including node turnover, node retention, and edge Jaccard similarity, were calculated across consecutive time points within each patient-tissue stratum.

### Size Normalization

Because many network metrics scale with repertoire size (node count), a normalization strategy was applied. For each metric, the observed value was regressed on log_10_(node count + 1) using either ordinary least-squares regression or a generalized additive model with splines (selected automatically based on a linearity test), and the residuals were retained as size-adjusted values (denoted with the suffix “_resid”).

### Predictive Modeling

We built binary classifiers to predict 3-month allograft outcome (Stable vs. Non-Stable, defined as borderline or reject combined) from pre-transplant (timepoint 0) PBMC network features. To avoid sequencing-depth artifacts, only size-normalized metrics, DRTC-based features, and the global clustering coefficient were used as candidate predictors; raw network size measures (node count, edge count, community count) were explicitly excluded. Feature count was constrained by the events-per-variable rule, permitting at most one feature per five minority-class events. Three algorithms were benchmarked: elastic net (α = 0.5) via glmnet (v4.1-10), random forest via ranger (v0.17.0, 500 trees, mtry = √p), and gradient-boosted trees via XGBoost (v3.1.2.1, 500 rounds). All models used leave-one-patient-out cross-validation with feature selection performed inside each fold to prevent information leakage. For each fold, features were ranked by absolute Wilcoxon test statistic on the training data and the top features (subject to the EPV constraint) were selected. Model discrimination was assessed by cross-validated AUC of the receiver operating characteristic curve (ROC) using pROC, and calibration by Brier score. The best-performing algorithm was selected based on AUC across the model comparison. Feature importance was assessed by selection frequency across cross-validation folds and by elastic net coefficient magnitude.

### Longitudinal Delta Analysis

To determine whether early post-transplant changes in network topology provide additional predictive information, per-patient deltas (3-month value minus pre-transplant value) were computed for all network metrics in both PBMC and Kidney specimens using paired samples at both timepoints. Univariate associations between delta values and binary outcome (Stable vs. Non-Stable) were tested using Wilcoxon rank-sum tests with Benjamini-Hochberg correction, and effect sizes were quantified by Cohen’s d. A weighted-average ensemble model was constructed by combining the baseline TCR prediction score (from the predictive modeling step) with a delta-based logistic regression submodel in PBMCs. The incremental contribution of delta features was evaluated by the DeLong test comparing ensemble and baseline-only AUCs, and by permutation testing (1,000 iterations) to obtain an empirical p-value for the delta model’s contribution.

### HLA Mismatch Analysis

Per-locus mismatches were counted as the number of donor alleles absent from the recipient genotype, evaluated across all donor-recipient allele combinations. Donor-recipient mismatch effects on network dynamics were assessed using linear mixed-effects models, with total mismatch count as a continuous predictor, log network size, timepoint, tissue, and clinical outcome as fixed effects, and patient as a random intercept. Tissue-stratified and tissue by HLA interaction models were also fitted. Locus-specific (HLA-A, -B, -DR), allele-specific, and MHC Class I versus Class II effects were examined separately. Longitudinal mismatch effects were assessed at 3, 6, and 12 months post-transplant using repeated measures mixed-effects models. To determine whether HLA features improve rejection prediction beyond TCR topology alone, the TCR-only model (from the predictive modeling step) were compared with TCR plus HLA combined models using DeLong tests on cross-validated AUCs and permutation testing as described above in the delta analysis.

### Statistical Analysis

All analyses was performed in R (v4.5.1) with visualizations using ggplot2 (v4.0.1) and pheatmap (v1.0.13) R packages. Associations between network features and clinical or biological factors were tested using linear mixed-effects models with the lme4 (v1.1-37) R package, incorporating a random intercept for the patient. Unless otherwise specified, fixed effects included status at 3 months, time point, tissue, and the centered log network size covariate. For each fitted model, ANOVA p-values, fixed-effect estimates, and marginal/conditional R² were reported using MuMIn (v1.48.11) R package. Multiple testing was controlled within each comparison and feature family using Benjamini-Hochberg FDR and Bonferroni corrections.

## Results

### Generation of Edit Distance Networks Across Sample Types

We previously published on a TCR repertoire cohort of 54 kidney transplant recipients classified at 3 months post-transplant as stable (n=33), borderline (n=15), or rejection (n=6).^11^ In total, 297 TCRβ sequencing runs were performed on serial peripheral blood and renal biopsy samples collected at pretransplant and at 3, 6, and 12 months post-transplant (Figure 1A). Patients received varied immunosuppression induction regimens, including alemtuzumab (campath, n=34), basiliximab (simulect, n=18), and methylprednisolone (solumedrol, n=2). When comparing unique clonotypes, defined as unique CDR3 amino acid and TCRβ V gene combinations, there were significantly more clonotypes identified per sequencing run in peripheral blood than in renal biopsies (*P* < 0.001, Figure 1B), consistent with the greater lymphocyte content of blood versus tissue. However, there was no significant difference in the number of clonotypes by graft status at 3 months, indicating that repertoire size alone does not distinguish outcomes (Figure 1C). To capture structural relationships within each repertoire, we constructed sequence-similarity networks for each sample, passing quality filters (minimum 100 clones and 1,000 reads). In these networks, nodes represent unique TCRβ CDR3 amino acid sequences and edges connect clonotypes that share the same V gene and have a normalized Levenshtein edit distance ≤ 0.3, where the edit distance is normalized by the mean CDR3 sequence length of the pair (Figure 1D). In total, 297 networks were constructed (183 PBMC, 114 kidney biopsy). Representative pretransplant networks, subsampled to equal clone counts to enable visual comparison, revealed apparent structural differences between tissue types and outcome groups, with rejection-associated networks appearing more fragmented and less densely connected than their stable counterparts (Figure 1E). To quantify these visual impressions systematically, we computed a panel of graph-theoretic metrics capturing connectivity, community structure, and global topology for each network. However, because network size is a known confounder of topological metrics, we first assessed the extent of this dependency across all features.

**Figure 1:**
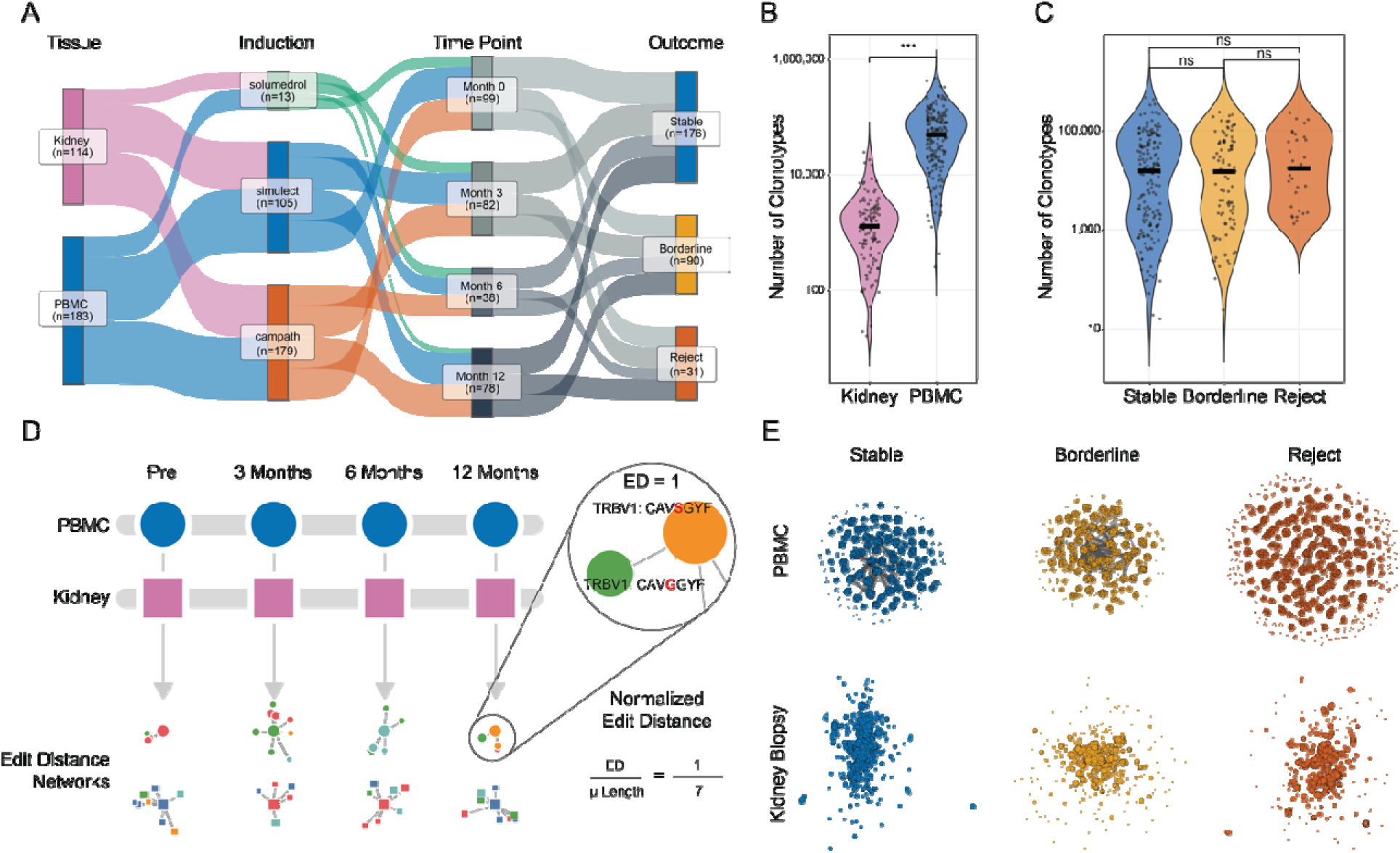
Application of edit-distance networks to a longitudinal TCRβ sequencing renal transplant cohort. **A**. Sankey diagram summarizing the cohort of 297 samples from 54 patients by tissue, immunosuppression induction, time point, and transplant outcome. **B**. Violin plot of the number of unique clonotypes per sequencing run by tissue. Central black bar indicates the mean. *** P < 0.001. **C**. Violin plot of the number of unique clonotypes per sequencing run by transplant outcome. Central black bar indicates the mean. ns, not significant. **D**. Schematic of normalized edit-distance-based network construction from paired tissue samples across longitudinal time points. Normalized edit distance is calculated as the edit distance (ED) divided by the mean CDR3 sequence length. **E**. Representative edit-distance-based networks from pre-transplant samples by tissue and transplant outcome. Node size is relative to the number of edges with each network subsampled to the equal number of clones to enable comparisons across groups.

### Size Correction is Required for Downstream Network Analysis

Network size, defined as the number of unique clonotypes, accounted for substantial variance across nearly all topological metrics (Figure 2A). This dependency spanned metrics with both strong positive and strong negative relationships to size, for example, degree entropy was nearly perfectly correlated with log-transformed node/clonotype count (Spearman *r* = 1.00), while density showed a strong inverse relationship (Spearman *r* = −0.93) (Figure 2B). Notably, the outcome groups (stable, borderline, reject) overlapped entirely along the size axis in both cases, reinforcing the need to correct for network size when comparing raw metrics across networks of different sizes. More complex, nonlinear relationships between size and individual metrics were also observed across the full panel of features (Supplemental Figure 1). To remove this confounding effect, each metric was modeled as a function of log_10_(node count) using both a linear and a generalized additive model (GAM), and the better-fitting model was selected for each metric. The resulting residuals were then extracted as size-corrected values for all subsequent analyses (Figure 2C). Post-correction, correlations between network size and all metrics were effectively eliminated (|*r*| < 10^-14^). A summary of the model selected (linear vs. GAM), along with the direction and magnitude of the size-metric correlation for each feature, is shown in Figure 2D. The residualization of the network metrics led to a substantial reduction in the observed tissue-specific differences in network topology (Supplemental Figure 2). All downstream analyses use these size-corrected residualized metrics.

**Figure 2.**
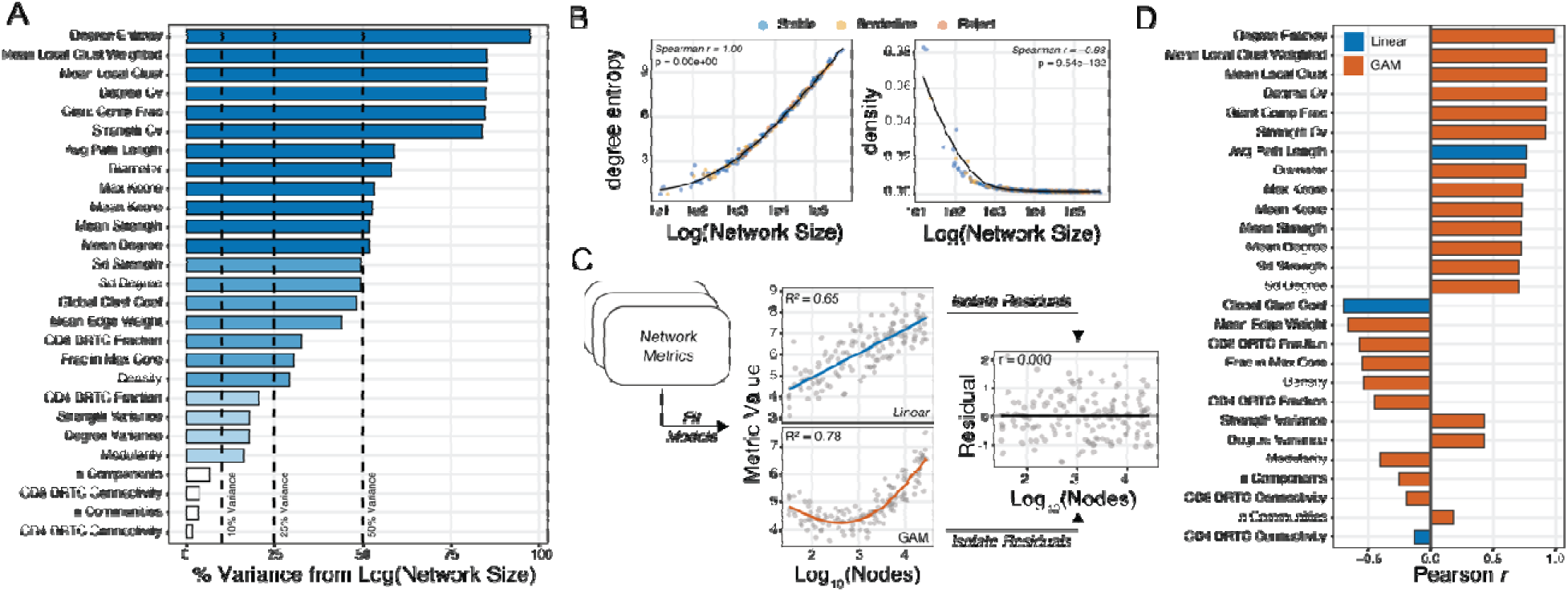
Network size drives substantial variance in network metrics, necessitating size-correction prior to downstream analysis. **A**. Percentage of variance in each network metric explained by network size (log-transformed), with metrics ranked in descending order. Dashed lines indicate 10%, 25%, and 50% variance thresholds. **B**. Representative scatter plots illustrating the relationship between network size and two network metrics, degree entropy (Spearman *r* = 1.00) and density (Spearman *r* = −0.93), with points colored by outcome classification (stable, borderline, reject). **C**. Schematic of the size-correction approach: a linear or GAM is fit to each metric as a function of log_10_(nodes), and the resulting residuals are extracted for subsequent analyses. **D**. Pearson correlation coefficients between each network metric and network size, shown separately for the linear (blue) and GAM (orange) models, illustrating the direction, magnitude, and model choice used for size correction.

### Graft Outcomes and Induction Therapy are Associated with Network Features

After establishing that size-corrected metrics are independent of repertoire depth, we next asked which biological and clinical factors drive variation in network topology in peripheral blood. For each residualized metric in PBMC samples, associations were tested against nine clinical variables: graft status at 3 months, induction therapy regimen, recipient age, and other transplant parameters. Effect sizes were quantified as η² from Kruskal-Wallis tests for categorical variables and ρ² from Spearman correlations for continuous variables (recipient age), while statistical significance was assessed using linear mixed-effects models with timepoint as a fixed-effect covariate and patient as a random intercept. The resulting bubble plot revealed that induction therapy regimen and graft status were associated with multiple topological features, while other clinical parameters showed more selective associations (PBMC, Figure 3A, Kidney, Supplemental Figure 3).

**Figure 3.**
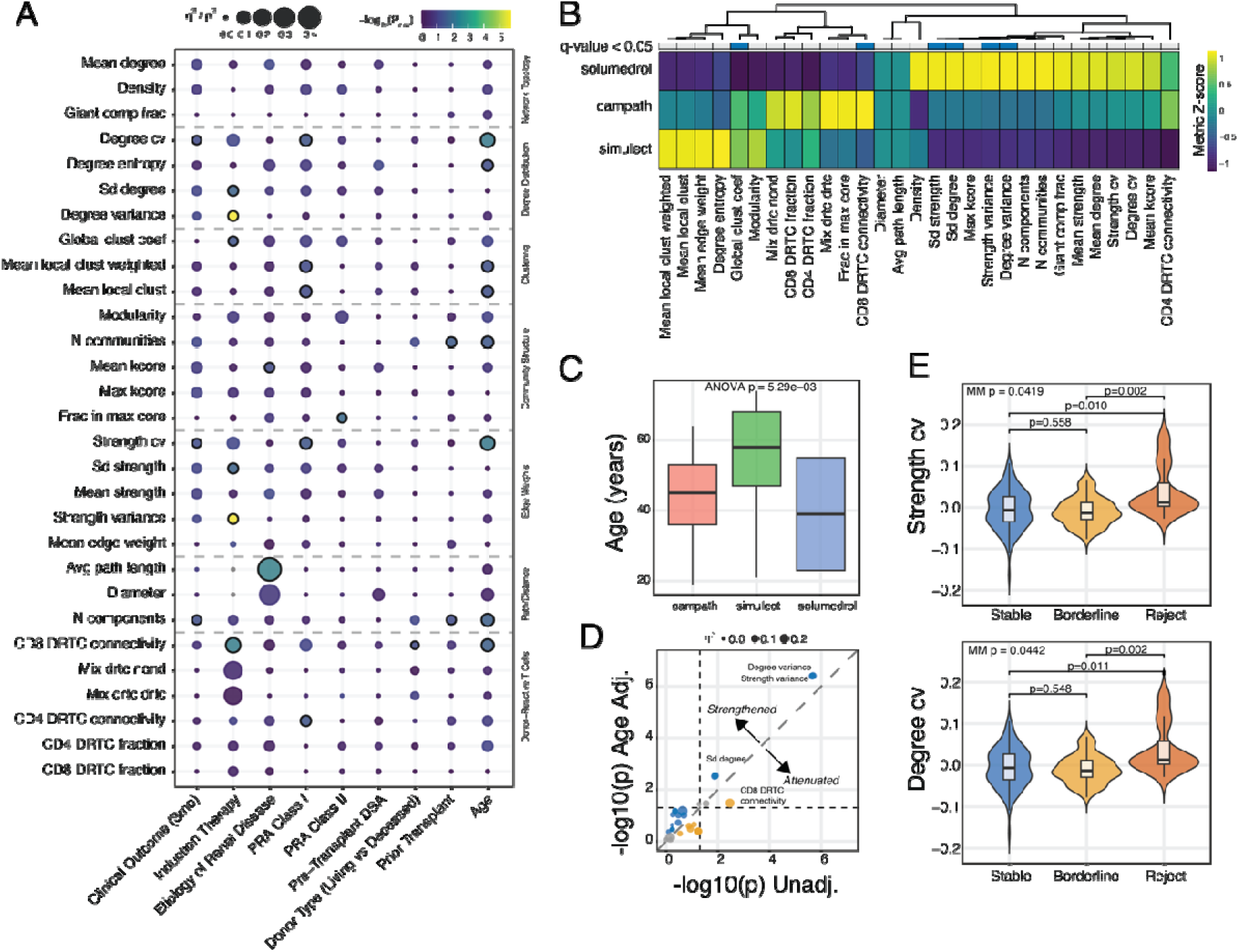
Network metric associations with clinical transplant parameters in peripheral blood. **A**. Bubble plot summarizing associations between nine clinical variables and size-corrected (residualized) TCR network metrics in PBMC samples. Bubble size encodes effect size (η² from Kruskal-Wallis for categorical variables; ρ² from Spearman correlation for age). Bubble color encodes statistical significance as −log_10_(raw mixed-effects model p-value), with timepoint and log-transformed repertoire size as fixed-effect covariates and patient as a random intercept. **B**. Summary heatmap of scaled group means across all residualized network metrics, stratified by induction therapy regimen (campath, simulect, solumedrol). Rows (groups) and columns (metrics) are hierarchically clustered. **C**. Boxplot of recipient age at transplant stratified by induction regimen. **D**. Scatterplot of the comparison of induction therapy associations before and after adjusting for recipient age. X-axis: unadjusted mixed-model effect sizes; y-axis: age-adjusted effect sizes. **E**. Violin plots of selected network metrics stratified by 3-month clinical outcome (stable, borderline, reject) in PBMC samples. Overlaid boxplots show medians and interquartile ranges. Overall p-values are from mixed-effects models (timepoint and log-transformed repertoire size as covariates, patient as a random effect). Pairwise comparisons use the Wilcoxon rank-sum test with the Benjamini-Hochberg correction.

The induction regimen emerged as a prominent driver of network variation, leading us to examine this association in greater detail. A summary heatmap of z-score-transformed residualized metrics, stratified by induction therapy (campath, simulect, solumedrol) and hierarchically clustered, revealed distinct topological profiles across regimens (Figure 3B). However, recipient age at transplant also differed across induction groups (Figure 3C), likely reflecting clinical practice patterns in which younger patients are preferentially selected for more potent lymphocyte-depleting agents. To determine whether the observed topology, induction associations reflect the immunosuppressive effect of the regimen itself or the underlying age-related immune characteristics of patients selected for each protocol, we compared unadjusted and age-adjusted mixed-model effect sizes (Figure 3D). After adjustment, several associations were strengthened (degree and strength variance) or attenuated (CD8 DRTC connectivity), suggesting that patient age at transplant partially accounts for the induction-related differences in network structure.

We next examined whether graft status at 3 months is associated with residualized network topology in PBMC. Violin plots of representative metrics confirmed outcome-associated differences at the individual-patient level: degree variance, strength variance, and other selected features differed across stable, borderline, and rejection groups (mixed-model P = 0.0419 and P = 0.0442 for degree and strength variance, respectively; Figure 3E). Post-hoc pairwise Wilcoxon rank-sum tests with Benjamini-Hochberg correction revealed significant differences between stable and rejection groups (P = 0.002) and between stable and borderline groups (P = 0.01), while borderline and rejection groups did not differ significantly (P > 0.5). This convergence between borderline and rejection groups suggests that network topology captures a graded continuum of immune perturbation rather than a discrete rejection-specific threshold. To resolve these subtle topological shifts into a more interpretable and consolidated feature space, we next applied principal component analysis (PCA).

### Network Connectivity and Structural Redundancy are Associated with Graft Outcome

To determine whether size-corrected network topology differs systematically by graft outcome, we performed PCA on the residualized metrics across all samples. The first two principal components accounted for 28.8% and 22.0% of the total variance, respectively, and revealed visual separation between outcome groups in the PCA space (Figure 4A). The top-loading features on PC1, degree entropy, mean degree, mean k-core number, and mean strength, all reflect network connectivity and structural redundancy (Figure 4B), suggesting that PC1 captures a coherent axis of repertoire organization ranging from densely connected to fragmented. To assess which clinical and demographic factors contribute to variation in network topology, we performed PERMANOVA on the PCA-derived coordinates. Graft status at 3 months and induction therapy each explained a significant proportion of the marginal variance (Figure 4C), confirming that an outcome-associated signal is present within these network metrics. When stratified by graft status at 3 months, PBMC-derived PC1 scores differed significantly across groups (Kruskal-Wallis P = 0.001, Figure 4D, upper panels) and were not observed to differ in renal samples (Figure 4D, lower panels). Patients who went on to experience borderline changes or rejection had lower PC1 scores, corresponding to more fragmented, less connected network architectures. Notably, this separation was driven by peripheral blood TCR repertoires, suggesting that features of the circulating T cell network topology detectable before surgery carry prognostic information. Given the clear separation along this composite axis, we next asked whether individual size-corrected network metrics could be combined into a formal predictive model of graft outcome.

**Figure 4.**
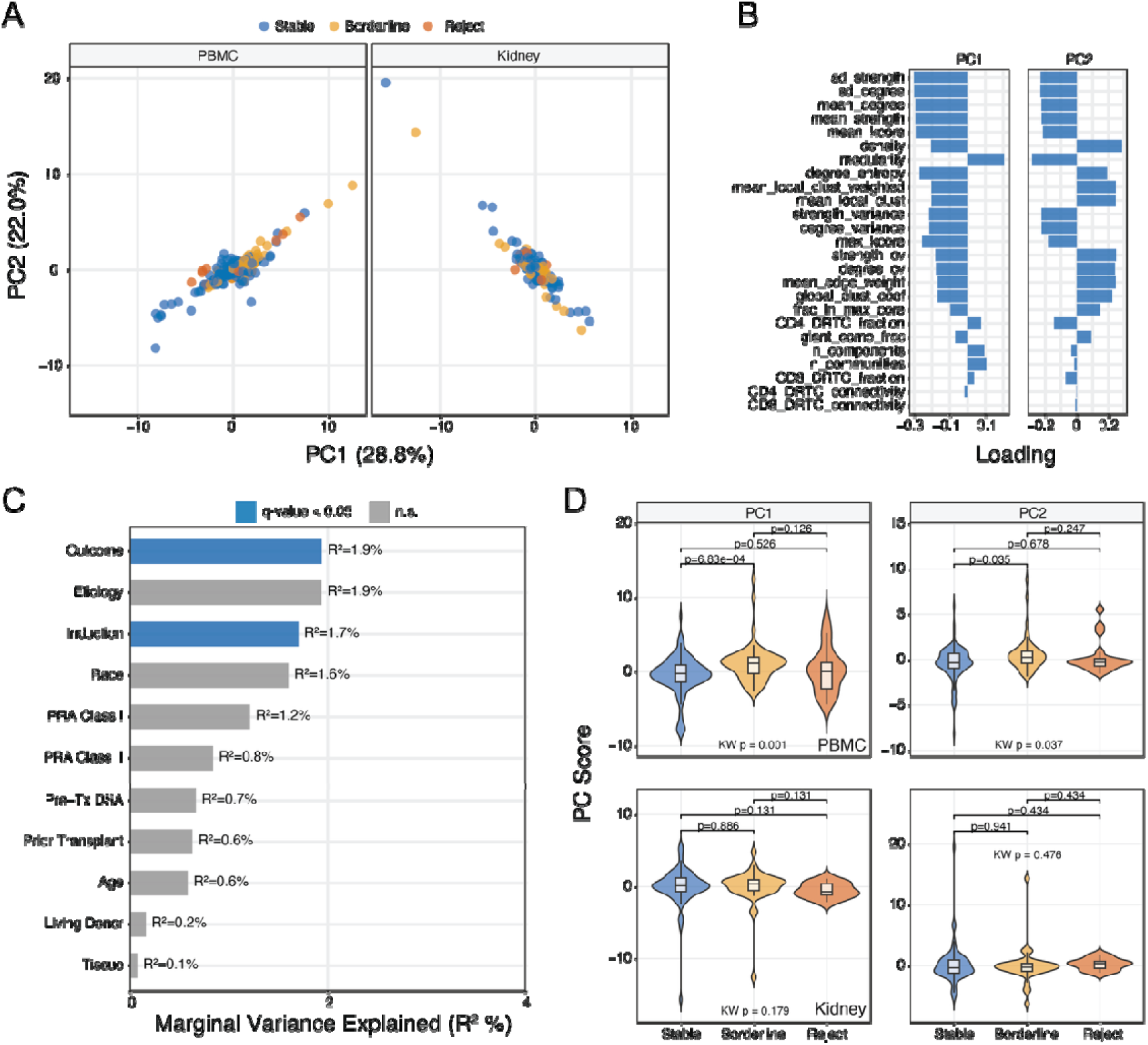
PCA of size-corrected network metrics reveals distinct profiles associated with graft outcome and induction therapy. **A**. PCA scatter plots of size-corrected network metrics for PBMC and kidney biopsy samples, with points colored by graft status at 3 months (stable, borderline, reject). **B**. PC1 and PC2 loadings for each network metric, ranked by magnitude of contribution. **C**. PERMANOVA analysis of marginal variance explained (R^2^) by clinical and demographic covariates across size-corrected network metrics. Bars represent the proportion of variance in PCA space attributable to each covariate, with significant associations (adjusted p ≤ 0.05) highlighted in blue and non-significant associations in gray. **D**. Violin plots of PC1 and PC2 scores for peripheral blood (upper panels) and kidney (lower panels) stratified by graft status at 3 months.

### Pre-Transplant PBMC Network Analyses May Predict Graft Outcome

Having established that size-corrected network topology differs by graft outcome, we asked whether these features could formally predict transplant outcome from a single pretransplant blood draw (Figure 5A). We assessed out-of-the-box performance of the network metrics across three model approaches: random forest, XGBoost, and elastic net models (Supplemental Figure 4). We trained the top-performing elastic net classifier (α = 0.5) on size-corrected network metrics derived exclusively from pretransplant PBMC samples, using a binary outcome of stable versus non-stable (borderline and rejection combined). To avoid information leakage, feature selection was performed independently within each fold of a leave-one-patient-out cross-validation (LOOCV) framework, and the maximum number of features was constrained by the events-per-variable rule (number of events / 20). The model achieved an area under the curve (AUC) of 0.808 (95% CI: 0.692–0.924), with a sensitivity of 76.2% and specificity of 75.8% at the optimal threshold determined by Youden’s index (Figure 5B). The classifier selected four features: degree entropy, mean degree, mean k-core number, and mean strength, with degree entropy contributing the largest coefficient to classification performance (Figure 5C). These are the same connectivity-related metrics that dominated PC1 in the exploratory analysis (Figure 4B), providing convergent evidence that network fragmentation is the key topological signature distinguishing outcome groups. Feature selection was stable across cross-validation folds (mean Jaccard index = 0.74), with each of the four core features selected in 72–98% of folds, indicating that the model is not driven by idiosyncratic fold-specific variable choices.

**Figure 5.**
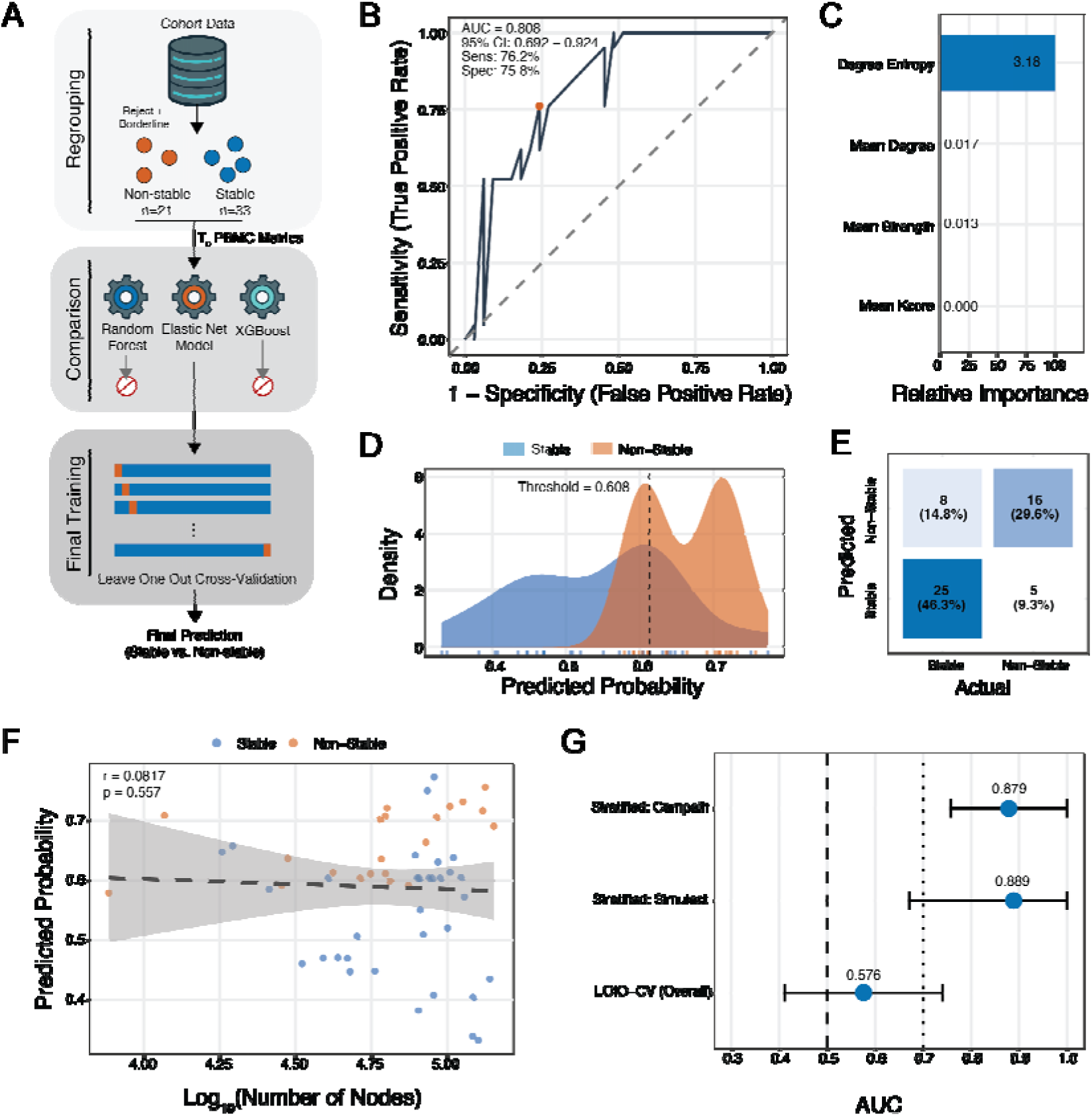
Predictive model based on pretransplant PBMC TCR profiling discriminates stable from non-stable kidney transplant outcomes. **A**. Schematic of the predictive modeling approach: samples were grouped into stable (n=33) or non-stable (n=21, including borderline and reject), the network topology metrics ability to predict were assessed by three model architectures, random forest, XGBoost, and elastic net. The top performing model was trained using leave one out cross-validation. **B**. ROC curve for the four-feature elastic net model trained on size-corrected network metrics, with an AUC of 0.808 (95% CI: 0.692–0.924), sensitivity of 76.2%, and specificity of 75.8%. The orange point indicates the optimal threshold. **C**. Relative importance of each feature in the elastic net model, with degree entropy contributing the most to classification performance. **D**. Density distributions of predicted probabilities for stable (blue) and non-stable (orange) outcomes, with the optimal classification threshold set at 0.608 (dashed line). **E**. Confusion matrix comparing predicted versus actual outcomes, showing overall classification accuracy with counts and percentages for each category. **F**. Scatter plot of predicted probability versus network size (log_10_ number of nodes), demonstrating no significant residual correlation between model predictions and network size (Spearman *r* = 0.0817, *p* = 0.557). **G**. Comparative AUC analysis showing performance of the overall model versus performance when stratified by induction therapy (campath and simulect) and leave-one-induction-out cross-validation (LOIO-CV), with point estimates and 95% confidence intervals. The dashed line indicates chance-level performance (AUC = 0.5) and the dotted line marks AUC = 0.7.

The predicted probability distributions for stable and non-stable patients showed clear separation at an optimal classification threshold of 0.608 (Figure 5D), and the corresponding confusion matrix confirmed balanced performance across both classes (Figure 5E). There was a clear bimodal distribution in the probabilities of non-stable outcomes (Figure 5D), this was not driven by a difference in borderline vs reject status or any other clinical parameter for the cohort (Supplemental Figure 5). Importantly, there was no residual correlation between predicted probabilities and network size (Spearman *r* = 0.0817, *P* = 0.557, Figure 5F), confirming that the size-correction procedure successfully removed this confounder and that the model is not simply recapitulating repertoire sampling depth. Finally, when performance was evaluated within each induction subgroup, using out-of-fold predictions from the full LOOCV model, filtered by induction type, the classifier maintained strong discrimination for both campath-treated (AUC = 0.879) and simulect-treated (AUC = 0.889) patients (Figure 5G). However, leave-one-induction- out cross-validation (LOIO-CV), in which the model was trained entirely on one induction group and tested on the other, yielded a substantially lower AUC of 0.576 (Figure 5G).

### Dynamic Changes in the TCR Repertoire, but not HLA Information, Improve Discriminative Prediction

The pretransplant model established that snapshot TCR network topology carries prognostic information, but transplantation initiates a dynamic immunological process in which peripheral and graft-resident T cell repertoires increasingly interact. To assess this, we first examined clonotype sharing between paired peripheral blood and kidney biopsy samples at multiple time points. The number of shared clonotypes between compartments increased over the first year post-transplant, with all outcome groups showing a shift toward greater blood–kidney overlap relative to pretransplant baselines (Figure 6A). While the overall Jaccard similarity index did not differ significantly between outcome groups at any single timepoint (Figure 6B, left), there was a significant temporal trend toward increasing clonotype sharing across the cohort (*P* = 0.0155, Figure 6B, right), consistent with progressive infiltration or equilibration of T cell clones between peripheral blood and the allograft. This increased overlap in PBMC and renal repertoires was more prominent at earlier stages for patients categorized as borderline or rejecting. This increased clonal exchange suggests that early post-transplant changes in the peripheral blood TCR repertoire may reflect active immunological engagement with the graft and could provide additional predictive information beyond a pretransplant snapshot alone.

**Figure 6.**
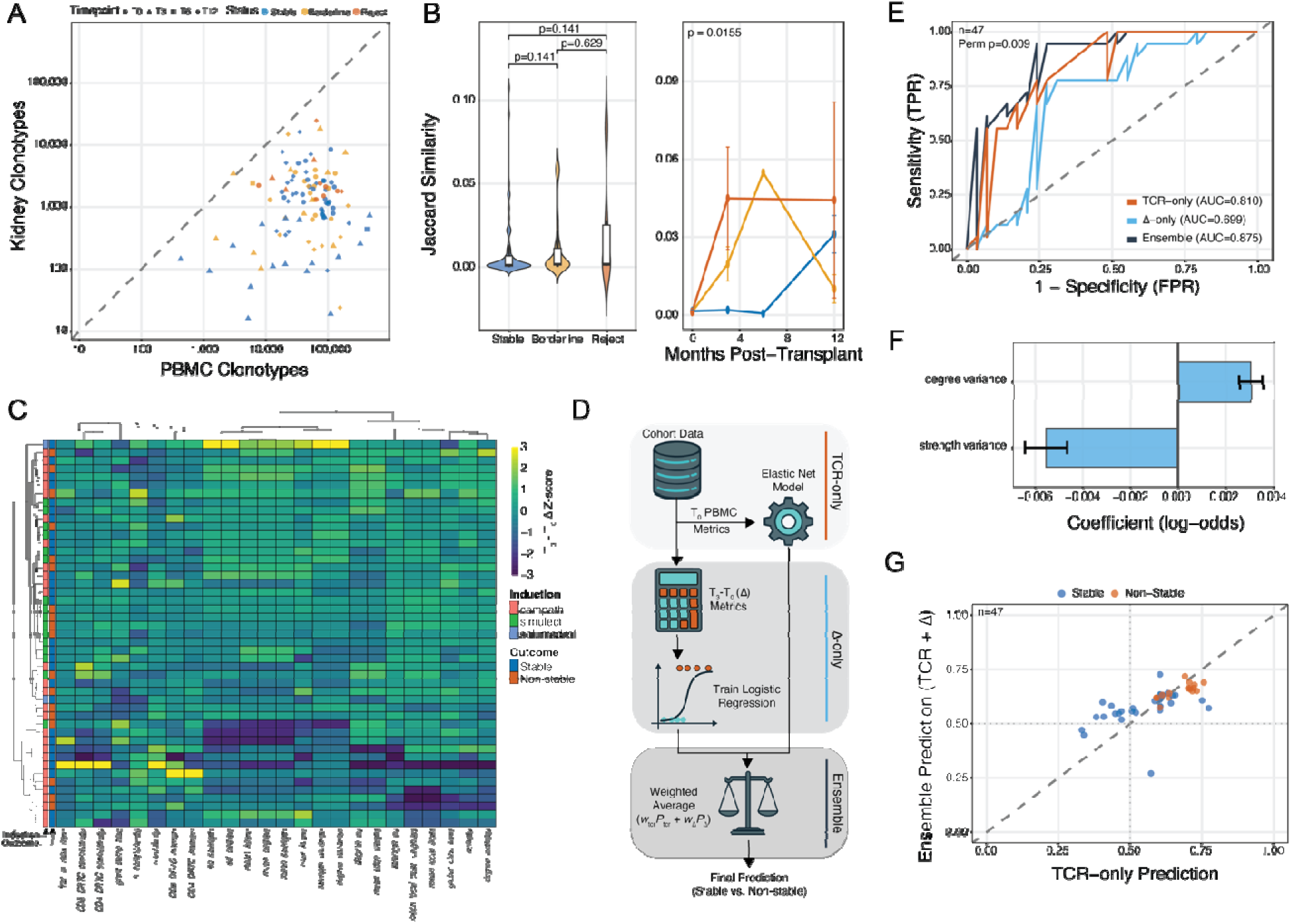
Longitudinal changes in network metrics improve prediction of graft outcome when ensembled with the baseline TCR network model. **A**. Scatter plot of shared clonotypes between PBMC and kidney biopsy, with points colored by graft status (stable, borderline, reject) and shaped by timepoint (T0, T3, T6, T12). The dashed line indicates equal clonotype counts across compartments. **B**. Violin plot of jaccard similarity index of shared clonotypes between peripheral blood and kidney across graft status groups (left) and line plot with longitudinal trend lines (right) showing temporal change of sharing (*p* = 0.0155). **C**. Heatmap of peripheral blood Δ network metrics (T_3_ − T_0_), with hierarchical clustering of patients and metrics. Rows are annotated by induction immunosuppression (campath, simulect, solumedrol) and graft outcome (stable, non-stable). **D**. Schematic of the ensemble modeling approach: the baseline elastic net model from Figure 5 (TCR-only) is combined with a logistic regression model trained on Δ metrics (Δ-only), and the two are integrated via a weighted average of predicted probabilities to generate a final stability prediction. **E**. ROC curves comparing the TCR-only (AUC = 0.810), Δ-only (AUC = 0.699), and ensemble (AUC = 0.875) models (n = 47), with the ensemble achieving the highest discriminative performance (permutation *p* = 0.009). **F**. Log-odds coefficients for the features selected in the Δ-only logistic regression model, showing opposing contributions of degree variance (positive) and strength variance (negative) to prediction of unstable outcome. **G**. Scatter plot comparing predicted probabilities from the TCR-only model (x-axis) versus the ensemble model (y-axis), with points colored by outcome (stable, non-stable).

To test this, we computed Δ network metrics (T_3_ − T_0_) for each size-corrected topological feature in peripheral blood and visualized these changes across patients by hierarchical clustering (Figure 6C). The resulting heatmap revealed partial separation of patients by both induction regimen and graft outcome, with non-stable patients tending to show distinct patterns of metric change in the first three months. To integrate the pretransplant and longitudinal signals, we constructed an ensemble model combining the static elastic net classifier from Figure 5 (TCR-only) with the Δ-only logistic regression via a weighted average of predicted probabilities (Figure 6D). The ensemble was evaluated on the 47 patients with paired pretransplant and 3-month PBMC samples. The TCR-only model maintained its performance in this subset (AUC = 0.810), while the Δ-only model provided a modest but independent signal (AUC = 0.699). The ensemble achieved the highest discriminative performance (AUC = 0.875; permutation *P* = 0.009), with optimal weighting of 60% and 40% for the Δ and TCR-only models, respectively (Figure 6E).

The Δ-only logistic regression model using the two features that survived events-per-variable-constrained selection: degree variance and strength variance (Figure 6F). These features showed opposing contributions; increasing degree variance (positive log-odds) and decreasing strength variance (negative log-odds) were both associated with non-stable outcomes. The improvement was not driven by a few reclassified patients; rather, the ensemble shifted predicted probabilities toward more confident and correct classifications across both outcome groups (Figure 6G). We applied the same ensemble framework using an HLA mismatch logistic regression in place of the Δ-only model, but adding HLA information to the TCR-only classifier yielded no improvement in discriminative performance (Supplemental Figure 6), indicating that the prognostic gain from the ensemble is specific to the longitudinal TCR network signal rather than conventional donor-recipient genotyping.

## Discussion

In this study, we demonstrate that the topological architectures of the pretransplant peripheral blood TCR repertoire can predict early kidney allograft outcomes from a single blood draw, achieving an AUC of 0.81 with 76% sensitivity and specificity. Incorporating early posttransplant network dynamics at three months further improved the discriminative performance to an AUC of 0.88. Critically, the predictive signal is independent of HLA mismatch and adding HLA mismatch information to the network-based model yielded zero improvement in performance (Supplemental Figure 6). These findings suggest that TCR repertoire profiling and network analysis may provide a different underlying framework for risk stratification. With the signal from TCR networks available before surgery, it may be clinically actionable for tailoring induction intensity or guiding the frequency of post-transplant monitoring.

The undertaking of this study was motivated by a methodological void in transplant repertoire studies. Nearly all of the current TCR and BCR literature within the field relies on discrimination using scalar diversity metrics, such as Shannon entropy, clonality indices, V-gene usage, or tracking individual clones.^5^ These metrics can be used to build sophisticated classification models^19^, but compress the relational architecture of the repertoire into a single number or limited vector space, discarding information of structurally related clones and clustering by sequence similarity. While some of these factors like V gene bias may appear to differ at baseline^6^, clonal expansion and diversity appear to be dynamic processes in the context of renal transplantation.^6,11,20,21^ Our finding that network fragmentation, low degree entropy, low mean degree, and reduced k-core structure predict non-stable grafts is a fundamentally different signal from “high clonality.” We propose that these represent different analytical windows onto the same underlying biology: scalar metrics detect clonal expansion, donor-reactive T cell profiling detects alloreactive clone abundance^10,11^, and network topology detects architectural fragmentation, three unique perspectives on alloreactive priming before transplantation. Biologically, high-frequency memory CD8⁺ donor-reactive T cells, which resist depletion across multiple induction agents^10,11^ likely serve as high-degree network hubs sharing sequence motifs with related clonotypes. In patients experiencing early rejection, dominance of a few such hubs with sparse surrounding connectivity would produce exactly the fragmented, low-entropy topology we observe. Previously, Sanders et al. explicitly called for “cluster and/or motif analyses” to elucidate rejection-associated epitopes.^11^ We directly address this gap with the creation of edit-distance networks in the current study using the same data set.

The independence of the network signal from HLA mismatch reflects a conceptual distinction between structural and functional risk. HLA mismatch quantifies the potential immunological distance between donor and recipient, a genotype-level measure of how different the graft appears to the host immune system. Network topology, by contrast, quantifies the current functional state of the recipient’s T cell compartment, how that immune system has been shaped by prior antigen exposures, homeostatic pressures, thymic selection, and immune history. In fact, TCR networks already encode a portion of the recipient HLA information and can be used to assign HLA genotyping for common HLA alleles.^22^ However, the clinical implication is significant: patients with identical HLA mismatch profiles may harbor very different network architectures and therefore substantially different rejection risks, a distinction invisible to current serological and genotypic screening. We note that the HLA analysis is limited by the smaller subset with complete typing (n = 37) and is based on only low-resolution typing, and while the results are internally consistent, they may be underpowered to detect modest additive effects.

In contrast, the Δ-enhanced ensemble model demonstrates that early post-transplant network remodeling provides prognostic information additive to the baseline snapshot, improving the AUC from 0.81 to 0.88, with optimal weighting favoring the Δ signal (60%) over the baseline (40%). This suggests that the trajectory of immune reorganization is more informative than the static pre-transplant state alone. The two selected delta features, degree variance and strength variance, showed opposing contributions: increasing degree variance and decreasing strength variance were both associated with non-stable outcomes. This pattern suggests that during the early post-transplant period, the degree distribution in non-stable patients becomes more heterogeneous, a few highly connected hubs emerging alongside many poorly connected nodes, while overall weighted connectivity becomes more uniform. These topological shifts are consistent with the selective persistence and expansion of CD8+ donor-reactive memory T cells despite induction therapy, a phenomenon now documented across both campath- and simulect-treated patients in this cohort^11^ and independently in rATG-treated recipients.^10^ As these resistant clones expand within an otherwise depleted or suppressed repertoire, they may generate the local connectivity hotspots that drive increasing degree variance while the loss of surrounding non-alloreactive clones homogenizes weighted strength.

A critical consideration in interpreting both the baseline and Δ models is the role of induction therapy as a source of structured heterogeneity. Campath induces profound lymphodepletion followed by reconstitution skewed toward memory T cells through homeostatic proliferation rather than thymopoiesis, resulting in a pruned yet alloreactively enriched repertoire in which CD8+ donor-reactive breadth paradoxically increases.^11^ In contrast, simulect largely preserves the pre-existing repertoire.^11^ Induction assignment in this cohort was not randomized; younger patients were preferentially selected for campath, reflecting clinical practice in which perceived immunological fitness guides induction intensity. Because age influences repertoire composition and network topology, the observed induction-associated differences are partially attributable to the baseline immune characteristics of patients selected for each protocol, as demonstrated by the attenuation of several associations after age adjustment (Figure 3C,D). This confounding by indication likely explains the strong within-induction prediction performance (Figure 5G, campath AUC = 0.879, simulect AUC = 0.889) but diminished cross-induction generalization (Figure 5G, LOIO-CV AUC = 0.576). Clinical implementation would therefore require protocol-aware calibration, similar to laboratory assays in which reference ranges are stratified by patient demographics and clinical context.

Several limitations should be considered when interpreting these results. First, the cohort of 54 patients from a single center, while sufficient for discovery, requires external validation. Second, the binary outcome classification (stable vs. non-stable) was necessitated by the small number of rejection events (n = 6); a three-class model distinguishing borderline from overt rejection would be clinically valuable but is underpowered in the current dataset. Third, cross-protocol generalization was limited, with LOIO-CV yielding an AUC of 0.58. We emphasize that this likely reflects genuinely different immunological processes rather than model failure: campath, basiliximab, and antithymocyte globulin create fundamentally different repertoire substrates^10,11^ Protocol-specific calibration, rather than a universal model, may be biologically appropriate. Fourth, the absence of phenotypic annotation (i.e., memory, naïve, effector, regulatory) of individual clones within networks could offer better insight into the functional clustering of the networks, linking network position to functional T cell states. This absence might be remedied using single-cell sequencing approaches that simultaneously capture TCR and gene/protein levels^23^, which may enhance predictive power and biological interpretability of the framework. These single-cell quantifications however, are at substantially lower yield in terms of TCR quantification (10^5^ compared to bulk 10^6–7^), but would include the TCR L chain that plays a role non-insignificant role in antigen recognition.^24^

Collectively, our findings establish TCR sequence-similarity network topology as a potential non-invasive, pre-transplant biomarker of kidney allograft rejection that captures functional immunological risk inaccessible to conventional HLA genotyping and serology. The convergence of evidence, 1) that network fragmentation reflects alloreactive priming detectable in a single blood draw, 2) that this signal is orthogonal to HLA mismatch, and 3) that early post-transplant remodeling provides additive prognostic value, support further exploration of TCR profiling in the context of solid organ transplant testing. Realizing this framework will require external validation in independent, multi-center cohorts to establish generalizability across sequencing platforms and demographic contexts, as well as protocol-stratified calibration to account for the distinct repertoire substrates created by different induction agents. Beyond transplantation, the analytical approach demonstrated here, rigorous size normalization of graph-theoretic features derived from CDR3 sequence-similarity networks, is broadly applicable to any setting in which TCR or BCR repertoire architecture carries clinical information, including autoimmunity, infection, and cancer immunology. Integration of donor-reactive T cell annotations into network structure, extension to BCR repertoires for antibody-mediated rejection, and pairing with single-cell phenotyping to link network position to functional T cell states represent immediate opportunities to deepen the biological interpretability of this framework. Prospective trials in which network-derived risk scores inform induction selection and monitoring intensity will ultimately determine whether TCR network analysis can transition from a discovery tool to a clinically actionable component of pre-transplant immunological assessment.

## Funding information

This study was funded by the Department of Defense through the Technology Development Award RT160073 (JL). JS was supported by the National Institute of Diabetes and Digestive and Kidney Diseases of the National Institutes of Health under Award Number T32DK077662 (Grant: T32DK077662/TR/ NIAID NIH HHS/United States).

## Declaration of Interest Statement

NB was previously employed by Santa Ana Bio, Inc and Omniscope, Inc and works as a scientific advisor to Epana Bio, Inc and as a consultant to Columbus Instruments. BB is a salaried employee and owns stock options in Adaptive Biotechnologies, Inc. The work presented does not pertain to any commercial endeavors in the companies listed above. The remaining authors do not have any pertinent disclosures.

## Abbreviations

AUC: Area under the curve
BCR: B cell receptor
CDR3: Complementarity-determining region 3
CFSE: Carboxyfluorescein diacetate succinimidyl ester
CI: Confidence interval
CMV: Cytomegalovirus
DRTC: Donor-reactive T cell clones
EBV: Epstein–Barr virus
EPV: Events-per-variable
FDR: False discovery rate
GAM: Generalized additive model
HLA: Human leukocyte antigen
LOIO-CV: Leave-one-induction-out cross-validation
LOOCV: Leave-one-patient-out cross-validation
MHC: Major histocompatibility complex
PBMC: Peripheral blood mononuclear cell
PCA: Principal component analysis
PERMANOVA: Permutational multivariate analysis of variance
rATG: Rabbit antithymocyte globulin
ROC: Receiver operating characteristic
TCR: T cell receptor
TCRβ: T cell receptor beta chain
V(D)J: Variable, Diversity, Joining (gene segments)
VDJdb: V(D)J database (curated TCR specificity database)

