## Supplemental Figures for "Pre-transplant TCR Network Topology Predicts Kidney Allograft Rejection Independent of HLA Mismatch"

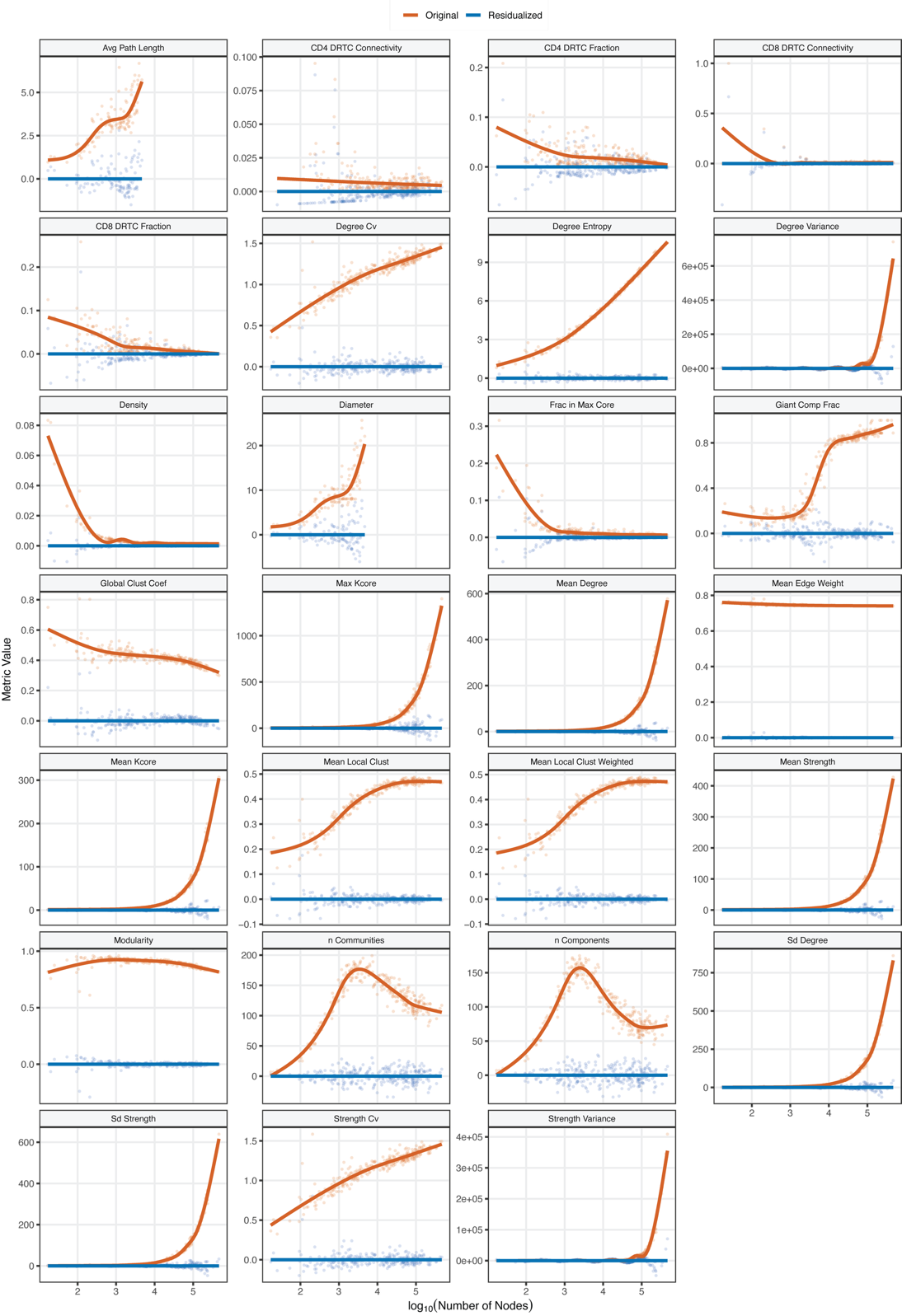

**Supplementary Figure 1:** Network size (x-axis, log10 transformed) and metric (y-axis) correlations for original (orange) and residualized (blue) values showing a reduction in size-dependent measures.

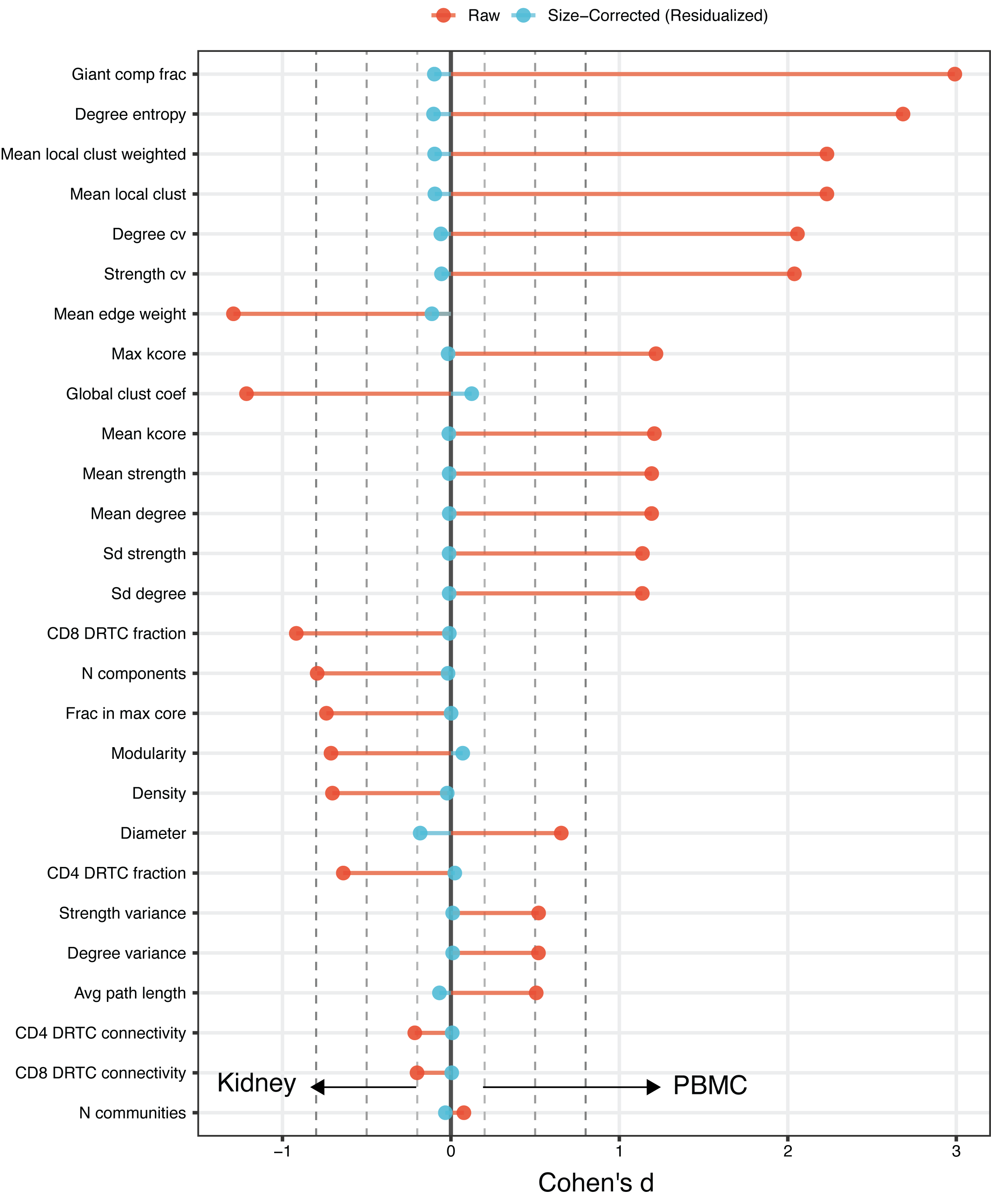

**Supplemental Figure 2**: Lollipop summary of the effect size (Cohen’s d) of network metrics comparing peripheral blood (right) to kidney biopsy (left) before (orange) and after (blue) size normalization.

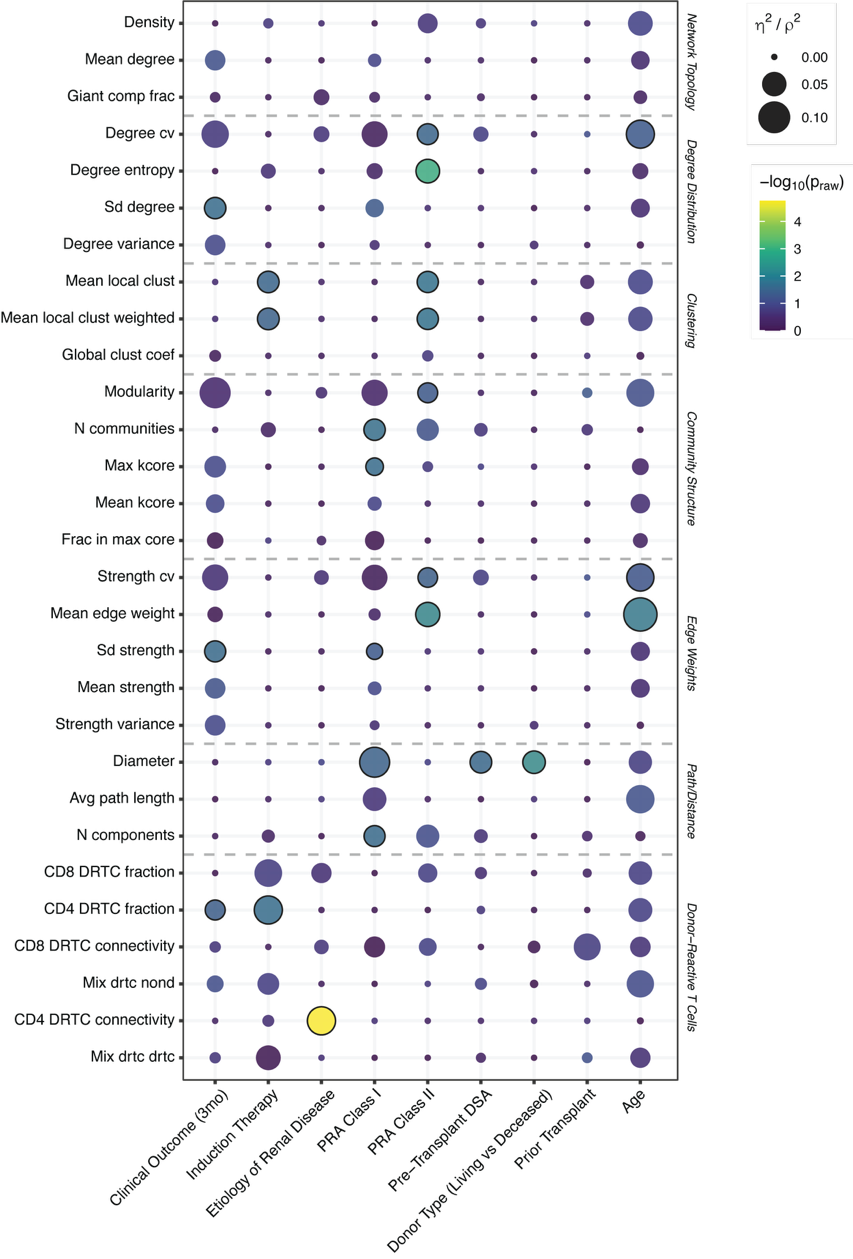

**Supplemental Figure 3**: Bubble plot summarizing associations between nine clinical variables and size-corrected (residualized) TCR network metrics in kidney samples. Bubble size demonstrates effect size (η² from Kruskal-Wallis for categorical variables; ρ² from Spearman correlation for age). Bubble color encodes statistical significance as −log_10_(raw mixed-effects model p-value), with timepoint and log-transformed repertoire size as fixed-effect covariates and patient as a random intercept.

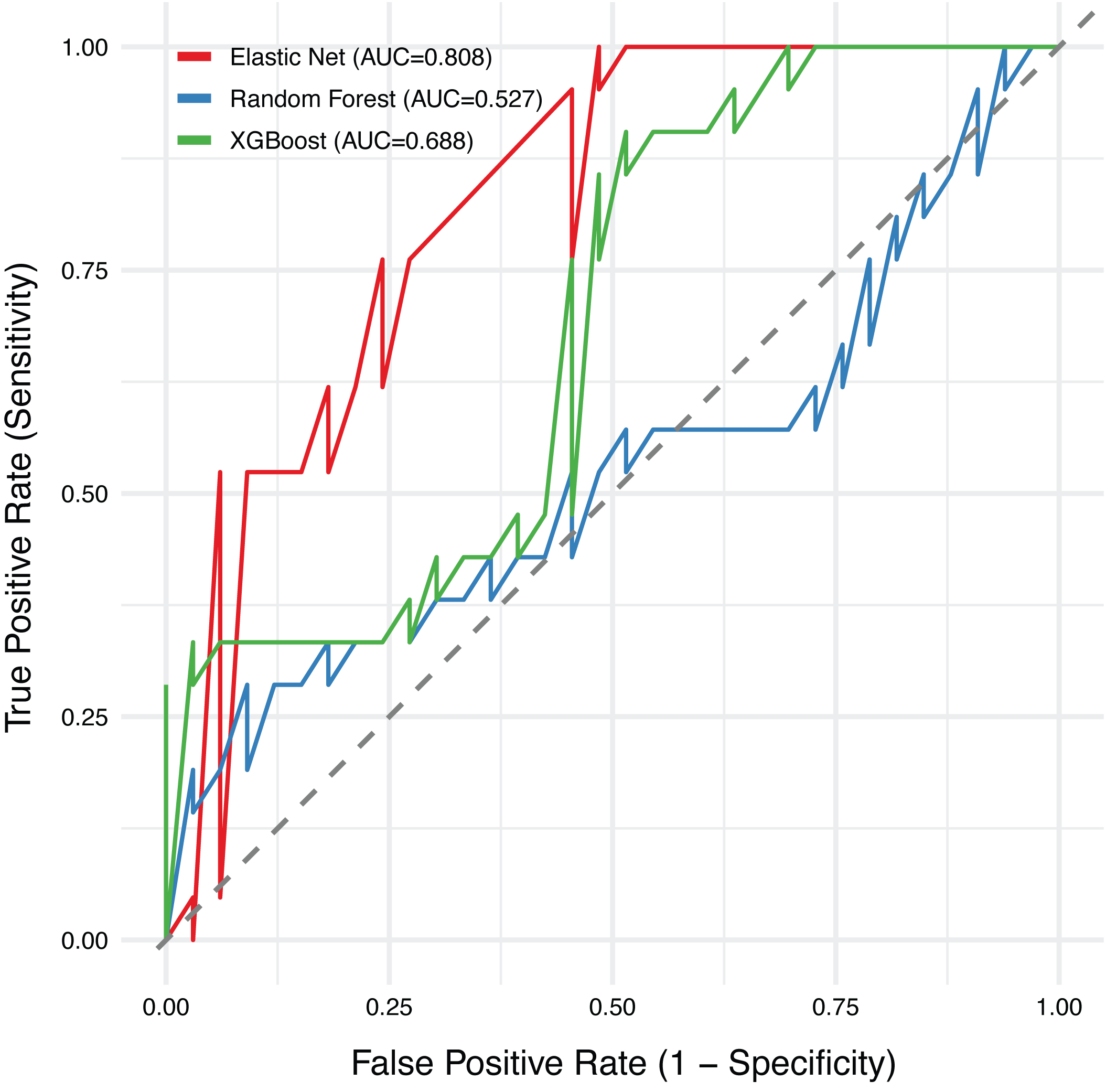

**Supplemental Figure 4**: ROC curves comparing out-of-the-box performance of the elastic net (red), random forest (blue), and XGBoost (green) models.

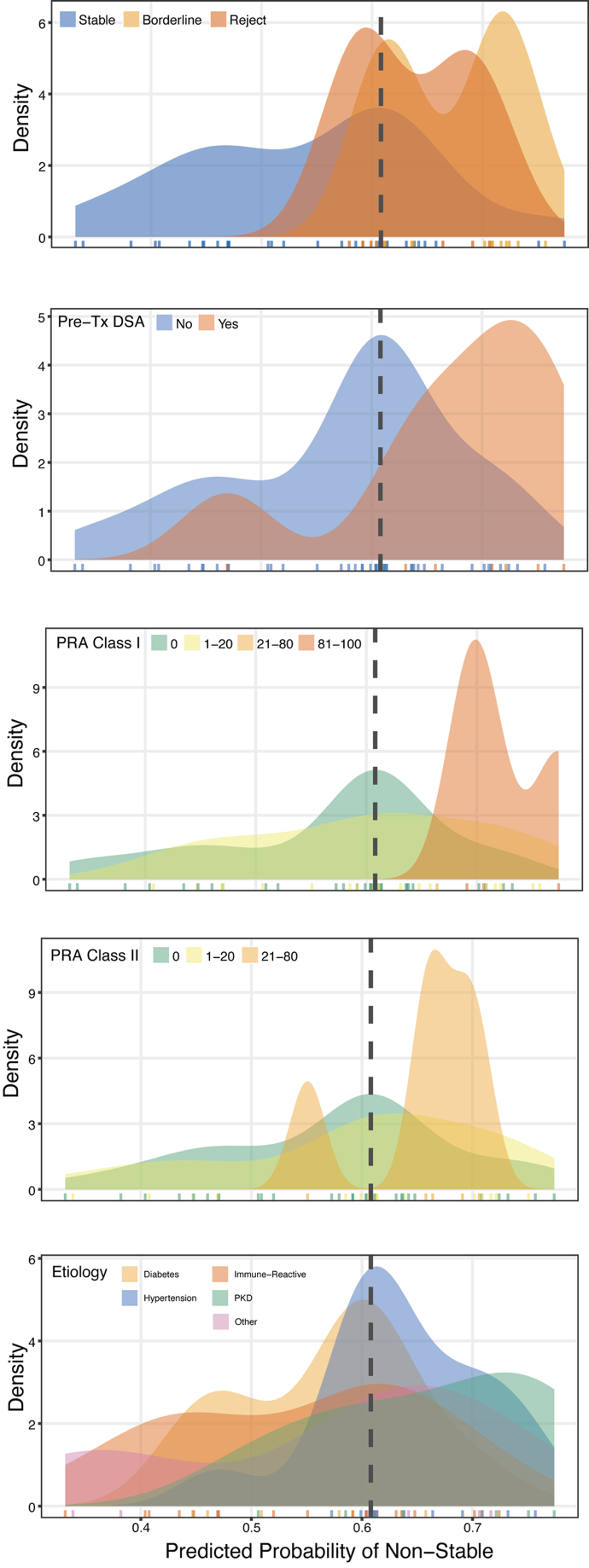

**Supplemental Figure 5**: Density distributions of predicted probabilities of TCR-only elastic net model by status at 3 months, previous transplant, PRA Class I, PRA Class II and underlying etiology of the end-stage renal disease.

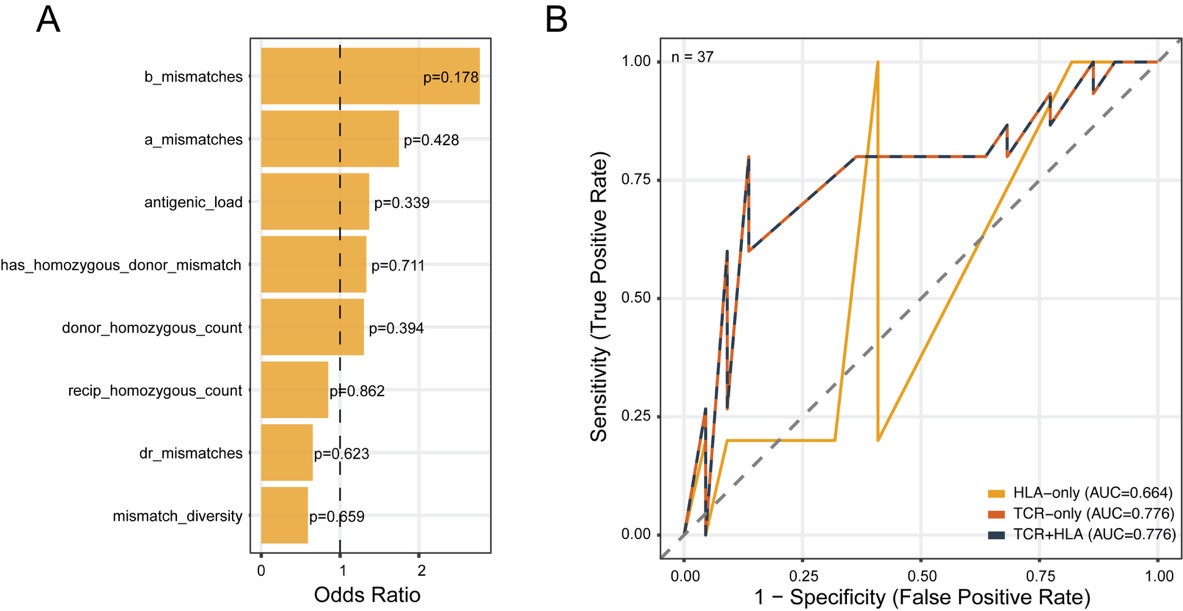

**Supplemental Figure 6**: **A**. Univariate log-odds ratio of non-stable outcomes per each HLA-based feature. **B**. ROC curves comparing the TCR-only (AUC = 0.776), HLA-only (AUC = 0.664), and ensemble (AUC = 0.776) models (n = 37).
